# A novel TIRAP-MyD88 inhibitor blocks TLR7 and TLR8-induced type I IFN responses

**DOI:** 10.1101/2025.04.25.650557

**Authors:** Kaja Elisabeth Nilsen, Jørgen Stenvik, Astrid Skjesol, Ingvild Bergdal Mestvedt, Siril Skaret Bakke, Miriam Soledad Giambelluca, Caroline S. Gravastrand, Kirusika Elamurugan, Liv Ryan, Terje Espevik, Maria Yurchenko

## Abstract

Endosomal toll-like receptors TLR7 and TLR8 are critical sensors of microbial RNA that initiate antiviral and antibacterial immune responses through type I interferon (IFN) and proinflammatory cytokine production. While TIRAP is traditionally associated with plasma membrane TLR signaling, recent evidence suggests it also contributes to signaling via endosomal TLRs. Here, we examined the role of TIRAP in TLR7/8 signaling using P7-Pen, a novel SLAMF1-derived peptide that disrupts the TIRAP–MyD88 interaction. In primary human monocytes and a whole blood model, P7-Pen inhibited TLR7- and TLR8-induced expression and secretion of IRF5-regulated cytokines IFNβ, IL-12p40, and IL-12p70, without effect on TNF or IL-6. Mechanistically, P7-Pen blocked TIRAP recruitment to the TLR8-MyD88 complex, leading to reduced late-stage IRAK1 activation, Akt and IKKα/β phosphorylation, and downstream IRF5 dimerization and nuclear translocation. Inhibition of *Staphylococcus aureus*-induced cytokine production by P7-Pen was associated with reduced bacterial phagocytosis, impairing endosomal delivery of bacterial RNA. Notably, P7-Pen failed to inhibit murine TLR7 responses, which correlated with a lack of TIRAP recruitment to MyD88 in mouse macrophages following TLR7 ligand stimulation, highlighting species-specific differences in TLR signaling mechanisms. These findings support a noncanonical role for TIRAP in regulating IRF5-dependent signaling downstream of human TLR7 and TLR8, and demonstrate that selective disruption of TIRAP recruitment by a SLAMF1-derived peptide effectively attenuates IFNβ production. This strategy may hold therapeutic potential in diseases characterized by dysregulated type I IFN responses, such as systemic lupus erythematosus and chronic infections.

## Introduction

Toll-like receptors (TLRs) are a family of pattern recognition receptors that play a central role in the innate immunity by detecting conserved molecular patterns associated with pathogens. TLRs are strategically positioned either on the cell surface or within endosomal compartments, enabling them to detect a broad spectrum of microbial components. These receptors can induce distinct immune responses, including the production of type I interferons (IFNs), proinflammatory cytokines, depending on the specific TLR and the context of activation (reviewed in (1, 2)). This diversity allows for a highly adaptable and effective immune response tailored to different classes of pathogens. Additionally, various regulatory molecules and pathways fine-tune the signaling of individual TLRs (reviewed in (2)). These mechanisms have been extensively studied in recent years to develop novel strategies for modulating TLR-driven inflammation in diverse pathological conditions.

A subset of TLRs—including TLR3, TLR7, TLR8, and TLR9—is localized in endosomal compartments, where they sense nucleic acids derived from certain viruses and bacteria. They play a critical role in the detection of intracellular as well as extracellular infections and mediate antiviral and antibacterial immune responses. Notably, there are species-specific differences in the expression and functionality of these receptors. In murine cells, both TLR7 and TLR8 are expressed; however, only TLR7 is responsive to ligand stimulation, and murine TLR8 is thought to be non-functional due to the absence of 5 amino acid motif required for RNA recognition (3). Human TLR7 and TLR8 sense single-stranded RNA (ssRNA), and have different ligand preferences with sequences rich in GU preferentially activating TLR7, and rich in AU sequences activate TLR8 (4). Also the expression of these TLRs varies between different subsets of immune cells (5).

TLR7 and TLR8 activation in endosomes triggers conformational changes that promote Toll/Interleukin-1 receptor (TIR) domains dimerization, enabling myeloid differentiation primary response gene 88 (MyD88) binding. This recruits interleukin-1 receptor-associated kinases 4 and 1 (IRAK4, IRAK1), forming the active Myddosome complex. The signal is further relayed through TNF-receptor-associated factor 6 (TRAF6) and transforming growth-factor-β-activated kinase 1 (TAK1), resulting in activation of mitogen-activated protein kinase (MAPK) pathways and inhibitor of nuclear-factor kappa B kinase subunit α/β (IKKα/β) heterodimer, which in turn activates nuclear-factor kappa B (NF-κB) and activator-protein 1 (AP-1) transcription factors (reviewed in (1, 6)). IKKβ also phosphorylates IRF5, which is required for IRF5 nuclear translocation (7, 8) and IRF5-induced transcription of *IFNβ* and *IL-12A* upon TLR8 activation (9).

Toll/interleukin 1 receptor domain-containing adaptor protein (TIRAP), a critical bridging adaptor for TLR2 and TLR4, was initially considered dispensable for signaling by endosomal Toll-like receptors (TLRs), including TLR7, TLR8, and TLR9, as supported by studies in murine cells for TLR7 and TLR9 (10, 11). However, more recent work by several groups, including our own, has revisited the role of TIRAP in modulating signaling downstream of endosomal TLRs, revealing that it may influence specific pathways (12–15). TIRAP has been shown to be critical for the assembly of the TLR9-Myddosome complex in murine iBMDMs (12). In plasmacytoid dendritic cells (pDCs), TIRAP knockout (KO) resulted in impaired production of IFNα in response to herpes simplex virus or influenza virus—sensed via TLR9 and TLR7, respectively—while IL-12p40 expression remained TIRAP-independent (12). Another study demonstrated that TIRAP KO reduced *Ifnb* expression in immortalized bone-marrow derived macrophages (iBMDMs) in response to R848 stimulation (ligand for murine TLR7), which was associated with decreased phosphorylation and activation of interferon regulatory factor 7 (IRF7) and extracellular signal-regulated kinase 1/2 (ERK1/2) (13). The role of TIRAP in regulating TLR7-mediated *IFNβ* expression in human cells has not yet been sufficiently explored.

Our previous study indicates that *TIRAP* silencing in human monocyte-derived macrophages (MDMs) does not alter TLR8-mediated ERK1/2 activation but instead leads to reduced phosphorylation of Akt kinase (14). TIRAP was recruited to the TLR8-Myddosome, which facilitated IRF5 nuclear translocation and *IFNB* and *IL12A* expression (14). Overall, based on previous findings, we suggest that targeting TIRAP recruitment to the TLR8-Myddosome—and potentially also the TLR7-Myddosome—may specifically inhibit TLR7/8-mediated expression of IRF5-dependent genes.

Recently, we developed and characterized a novel SLAMF1-derived peptide, P7, conjugated to the cell-penetrating peptide penetratin (Pen) for intracellular delivery (P7-Pen) (16). The P7-Pen peptide disrupts several key protein-protein interactions (PPIs) in human cells, including TRAM–SLAMF1, TRAM–Rab11FIP2, and TIRAP–MyD88 (16, 17). By targeting these PPIs, P7-Pen effectively inhibits TLR4-mediated proinflammatory cytokine expression and secretion in human monocytes and macrophages, prevents mortality in a murine LPS-shock model and improves heart function in myocardial infarction model (16, 18). Given its ability to block the TIRAP–MyD88 interaction, we hypothesized that P7-Pen might also inhibit the IRF5-dependent axis of TLR7/8 signaling.

In this study, we investigated whether the P7 peptide can inhibit TLR7- and TLR8-mediated signaling by blocking TIRAP recruitment to the Myddosome, and whether this inhibition regulates signaling mechanisms such as Akt phosphorylation. Using primary human monocytes, a human whole blood model, and a panel of TLR7/8 ligands, we found that P7-Pen effectively suppressed the expression and secretion of IRF5-dependent cytokines—including IFNβ, IL-12p40, and IL-12p70—downstream of TLR7 and TLR8 activation. Additionally, we investigated the molecular mechanisms of endosomal TLR signaling regulation by P7-Pen in human cells, and further examined those in primary murine macrophages.

## Materials and methods

### Primary cells, cell lines

Human buffy coats and serum were from the blood bank at St. Olavs Hospital (Trondheim, Norway), with approval by the Regional Committee for Medical and Health Research Ethics (REC) in Central Norway (no. 2009/2245). Primary human monocytes were isolated from the buffy coat by adherence, as previously described (16). After isolation, monocytes were kept overnight in RPMI1640 (Sigma, Merck, Darmstadt, Germany), supplemented with 30% of pooled human serum, followed by media change to RPMI1640 with 10% human serum prior experiment. THP-1 monocytic cell line derived from acute monocytic leukemia (ATCC TIB-202) were cultured in RMPI 1640 supplemented by 10% heat-inactivated FCS, 100 U/ml penicillin, 100 μg/ml streptomycin (pen/strep) (Thermo Fisher Scientific), and 5 μM β-mercaptoethanol (Sigma-Aldrich, Merck). Prior experimental procedures THP-1 cells were differentiated with 60 ng/ml of phorbol 12-myristate 13-acetate (PMA) (Sigma-Aldrich, Merck) for 48 h, followed by 48 h in medium without PMA. Media were changed to fresh media before the pretreatments and stimulation. The iBMDMs (immortalized bone-derived-macrophages) from wild type C57BL/6 mice were made in the lab of Dr. Douglas T. Golenbock (19).

### Bone marrow-derived immature DCs (iDCs) and macrophages (BMDMs)

BMDMs and iDCs were generated from BM cells from 10- to 12-wk-old C57BL/6 mice. Briefly, BM cells were flushed out from the femurs. After lysis of RBC, whole BM cells (2 × 10^5^ cells/ml) were cultured in 100-mm^2^ culture dishes in 10 ml/dish RMPI 1640 medium supplemented by 10% FCS and antibiotics (pen/strep). For generation of iDCs media was supplemented by 20 ng/ml mGM-CSF and 20 ng/ml mIL-4 (Sigma, Merck), and for generation of BMDMs – 10 ng/ml mM-CSF (Sigma, Merck). At day 2, another 5 ml of fresh complete medium containing cytokines was added. On day 4 of the culture, half of the medium was carefully changed. On day 6, non-adherent DCs and loosely adherent DCs were harvested by gently pipetting and used as iDC. iDCs recovered from these cultures were ∼60% CD11c^+^, ∼85% CD80^+^, ∼36% CD86^+^, and ∼2% CD40^+^ as analyzed by flow cytometry using BD LRSII flow cytometer, the FACS Diva software (BD Biosciences, NJ, USA). Data were exported and analyzed with FlowJo software v10.0.5 (Tree Star, Ashland, OR, USA).

#### Reagents

Synthetic peptides for assays with living cells were from Thermo Fisher Scientific, with sequences RQIKIWFQNRRMKWKK for Penetratin (Pen), ITVYASVTLTG-Pen for P7-Pen, IATYASTALTG-Pen for C3-Pen control peptide. Peptide modifications and characteristics: N-terminal acetylation, C-terminal amidation, >90% purity, guaranteed TFA removal, control for endotoxin levels (less then <10 EU/mg). Ultrapure K12 LPS and B4 LPS (0111:B4) from *E. coli*, ultrapure thiazoloquinoline compound CL075, imidazoquinoline compounds R837 (Imiquimod), R848 (Resiquimod), benzazepine analog TL8-506 were from InvivoGen (San Diego, CA, USA). For stimulation of the primary cells, LPS was used at concentration 100 ng/mL, TLR7/8 ligands – 1 µg/mL if not indicated otherwise in the figure legends. Selective allosteric pan-Akt inhibitor Miransertib (#1313881-70-7, Med-ChemExpress) was diluted in DMSO at concentration 5 mM and stored at −80 °C; working solutions (2 μM) were prepared in cell-culture media immediately before use.

#### Antibodies

The following primary antibodies were used: mouse GAPDH (ab9484), rabbit β-tubulin (ab6046) from Abcam (Cambridge, UK); mouse β-tubulin (D3U1W, #86298), rabbit phospho-Akt (Ser473) (D9E, #4060), phospho-p38 MAPK (T180/Y182), phospho-STAT1 (Tyr701) (58D6), phospho-TAK1 (T184/187) (90C7), IRAK1 (D51G7), MyD88 (D80F5), phospho-IKKα/β (Ser176/180) (16A6, #2697) from Cell Signaling Technology (Danvers, MA, USA); rabbit PCNA Abs were from Santa Cruz Biotech (Santa Cruz, CA, USA); sheep IRF5 and IRAK4 were from MRC-PPU Reagents (University of Dundee, Dundee, United King-domUK); goat TIRAP polyclonal Abs were from Invitrogen (#PA5-18439, Waltham, MA, USA). Secondary antibodies (HRP-linked) were from DAKO Denmark A/S (Glostrup, Denmark).

#### RT-qPCR

Total RNA was isolated from the cells using Qiazol reagent (QIAGEN, Germantown, AR, USA), and chloroform extraction was followed by purification on RNeasy Mini columns with DNAse digestion step (QIAGEN). cDNA was prepared with a Maxima First Strand cDNA Synthesis Kit for a quantitative real-time polymerase-chain reaction (RT-qPCR) (ThermoFisher Scientific, Waltham, MA, USA), in accordance with the protocol of the manufacturer, from 400–600 ng of total RNA per sample. Q-PCR was performed using the PerfeCTa qPCR FastMix (Quanta Biosciences, Gaithersburg, MD, USA) in repli-cates and cycled in a StepOnePlus™ Real-Time PCR cycler (ThermoFisher Scientific, Waltham, MA, USA). The following TaqMan® Gene Expression Assays (Applied Biosys-tems®, ThermoFisher Scientific, Waltham, MA, USA) were used: *IFNβ* (Hs01077958_s1), *TNF* (Hs00174128_m1), *TBP* (Hs00427620_m1), *IL-6* (Hs00985639_m1), *IL-1β* (Hs01555410_m1) for human cells; *Ifnβ* (Mm00439552_s1), *Tnf* (Mm00443258_m1), *Il-1β* (Mm00434228_m1), and *Tbp* (Mm01277042_m1) for murine cells. The level of *TBP/Tbp* mRNA was used for normalization and the results presented as a relative expression compared to the control’s untreated sample. Relative expression was calculated using Pfaffl’s mathematical model. Graphs and statistical analyses were made with GraphPad Prism v10.1.2 (Dotmatics, Bishops Stortford, UK), with additional details provided in the figure legends or statistics paragraph.

#### Whole blood assay

Blood samples were obtained from healthy volunteers that gave signed consent for experimental procedures, approved by REC in Central Norway (REK# S-04114). Refludan (lepirudin) was used as anticoagulant in 50 µg/ml concentration as described before (20). Blood samples were distributed to sterile polypropylene tubes, 0.25 ml per sample, with total volume of reagents added to each sample being 0.1 ml (in PBS). After addition of 20 μM peptide or solvent (water, H2O) for 30 min, samples were stimulated with LPS (100 ng/ml), CL075, R837 (1 μg/ml) or 2 x10^6^ of pHrodo red *E. coli* or 4x10^6^ of pHrodo red *S. aureus* bioparticles for 5 h at 37°C (slow rotation). Blood was transferred to tubes containing ethylenediamine tetra acetic acid (EDTA, 10 mM), spun down at 3220 × g at 4°C for 15 min for plasma separation, and the plasma frozen for later cytokine expression analysis.

#### ELISA, BioPlex and LDH Assays

IFNβ levels were determined using assays from PBL Assay Science (Piscataway, NJ, USA): VeriKine-HSTM Human Interferon-Beta Serum ELISA Kit (#41415, PBL Assay Science). Other cytokines for human primary cells were analyzed using BioPlex cytokine assays from Bio-Rad, in accordance with the instructions of the manufacturer with the recommended concentration of reagents, but in reduced volume (1:2), using the Bio-Plex Pro™ Reagent Kit III and Bio-Plex™ 200 System (Bio-Rad, Hercules, CA, USA). TNF, IL-6 and IL-8 ELISA were used instead of Bio-Plex when indicated in the figure legends and were human TNF-alpha DuoSet ELISA (DY210), IL-6 DuoSet ELISA (DY206) and IL-8/CXCL8 Duoset ELISA (DY208) (R&D Systems, Minneapolis, MN, USA). CyQUANT LDH Cytotoxicity Assay (Thermo Fisher Scientific, Norway) was used to measure the extracellular LDH in supernatants as suggested by manufacturer.

#### Phagocytosis flow cytometry assay

A flow cytometry-based phagocytic assay was used to measure the phagocytic efficiency of red pHrodo-conjugated BioParticles in primary human monocytes. Following reagents were used: pHrodo Red *E. coli* K12 BioParticles (P35361), pHrodo Red *S. aureus* BioParticles (A10010) from Invitrogen. Prior to being added to cells the bacterial bioparticles were diluted in stimulation media. After stimulation, cells in 6-well plates were placed on ice, washed with cold PBS, detached by treatment with Accutase solution for 10–15 min (Sigma, Merck, Darmstadt, Germany) and transferred into FACS tubes. The cells were washed with PBS containing 2% FCS, followed by PBS. The fluorescence intensity was measured with a BD LRSII flow cytometer using the FACS Diva software (BD Biosciences, NJ, USA). Data were exported and analyzed with FlowJo software v10.0.5 (Tree Star, Ashland, OR, USA).

#### Live bacteria uptake

THP-1 cells were plated at 2 × 10^5^ cells/well in 24-well plates and differentiated with PMA as described in the Primary cells and cell lines section for 5 d (without antibiotics in media). Adherent differentiated cells have got fresh RPMI 1640 with 10% FCS. Live *E*. *coli* (DH5α) and *S*. *aureus* (protein A negative strain Wood 46) were grown to optical density of 0.35 at 600 nm, washed and diluted in PBS and given at a dose of 25 bacteria per cell in 4-5 replicates. Bacteria were centrifuged onto differentiated THP-1 monolayers at 750 g for 7 min at 4°C. Plates were warmed up to 37°C in a water bath for 15 min, and quickly transferred to ice. Each well was then washed 3× with ice-cold PBS and incubated with warm 10% FCS RPMI medium containing 100 μg/ml of gentamycin for 30 min at 37°C to remove extracellular bacteria. Wells were washed 2× by cold PBS and lysed in 1 ml sterile water. Viable bacteria counts were determined by plating 10 μl of 10-fold dilutions, onto Luria-Bertani agar (in triplicates, to account for technical pipetting error), and plates incubated at 37 °C for 16 h. Colony-forming units (CFU) were counted and the number of bacteria per cell calculated.

#### Immunoprecipitations

PBMC-derived monocytes for endogenous IPs were lysed using 1 X lysis buffer (150 mM NaCl, 50 mM TrisHCl (pH 8.0), 1 mM EDTA, 0.5% NP40) and supplemented with EDTA-free Complete Mini protease Inhibitor Cocktail Tablets as well as a PhosSTOP phosphatase-inhibitor cocktail from Roche, with 50 mM NaF and 2 mM Na_3_VO_3_ (Sigma, Merck, Darmstadt, GermanySigma). Immunoprecipitations (IPs) were carried out on rotator at +4 ℃ overnight by co-incubation of the lysates from the stimulated cells (500 μg of protein/IP) with specific anti-TIRAP antibodies covalently coupled to Dynabeads (M-270 Epoxy, Thermo Fisher Scientific, Waltham, MA, USA). IPs were washed by lysis buffer four times, followed by elution of co-precipitated complexes by heating the samples in a 1×loading buffer (LDS, Invitrogen) without reducing reagent to minimize the antibodies’ leakage to the eluates. Eluates were transferred to clean tubes, followed by the addition of reducing reagent dithiothreitol (DTT) (Sigma, Merck, Darmstadt, Germany) to the 40 mM concentration, samples were heated and analyzed by SDS-PAGE and WB. Precipitates were loaded to the gels in parallel with respective whole cell lysates (WCLs) for input control.

#### Western Blotting and SDS-PAGE

Cell lysates for pSTAT1 analysis were prepared by simultaneous extraction of proteins and total RNA using Qiazol reagent (QIAGEN, Germantown, AR, USA), as suggested by the manufacturer. Protein pellets were dissolved by heating the samples for 10 min at 95 °C in a buffer containing 4 M urea, 1% SDS (Sigma, Merck, Darmstadt, GermanySigma), and NuPAGE® LDS Sample Buffer (4X) (Thermo Fisher Scientific, Waltham, MA, USA), with a final 25 mM DTT in the samples. Otherwise, lysates were made using 1X RIPA lysis buffer with inhibitors of proteases and phosphatases (16). Samples were loaded to precast NuPAGE™ Novex™ protein gels (ThermoFisher Scientific). Proteins on gels were transferred to iBlot Transfer Stacks by using the iBlot Gel Transfer Device (ThermoFisher Scientific). The blots were developed with the SuperSignal West Femto (ThermoFisher Scientific) and visualized with the LI-COR ODYSSEY Fc Imaging System (LI-COR Biotechnology, Lincoln, NE, USA). For densitometry analysis of the bands, Odyssey Image Analysis Studio 5.2 software (LI-COR Biotechnology, Lincoln, NE, USA) was used, and the relative numbers of bands’ intensity were normalized to the intensities of the respective loading-control protein (GAPDH, or PCNA, or β-tubulin). Loading-control protein expression was always performed on the same membrane as the protein of interest.

#### Native gel for IRF5 dimerization

To study the dimerization of IRF5, we used a modified protocol published by Lopez-Pelaez et al. (7). For this, 8% native gels (Novex Tirs-Glycin Plus Wedge Well, Invitrogen) were prerun for 30 min at 40 mA at +4 °C in 25 mM Tris-192 mM glycine pH 8.4 with 0.1% (wt/vol) deoxycholate (DOC) in the cathode chamber only. Cell extract (15 μg protein) containing 1% (wt/vol) DOC was diluted with 2 x native gel buffer (Novex Native Tris-Glycine sample buffer) and electrophoresed at +4 °C for 60 min at 25 mA. The gels were incubated for 30 min at RT with 25 mM Tris-192 mM glycine pH 8.4, 0.1% (wt/vol) SDS, transferred to nitrocellulose iBlot Transfer Stacks, and immunoblotted with the IRF5 antibodies and secondary antibodies as described in Western Blotting section.

#### Immunofluorescence and Scan^R Analysis

Monocytes were isolated from PBMC using CD14 MicroBeads UltraPure (Miltenyi Biotech) and seeded in 96-well glass-bottom plates (P96-1.5H-N, Cellvis, CA, USA) at 50000 cells/well and in 24-well culture plates (250000 cells/well). Cells were left untreated or stimulated with CL075 (2 μg/mL) for 60 min, with four technical replicates per condition. Fixation, immunostaining with anti-human IRF5 mAb (Abcam, #10T1) and anti-human p65 XP mAb (Cell Signaling Technology, #8242, Danvers, MA, USA), and Scan^R high-throughput imaging (Olympus Europa SE & CoOlympus, Hamburg, Germany) were done, as previously described in detail (9). Quantification of IRF5 nuclear translocation was done with Scan^R analysis software (v2.8.1) and calculated as a percentage of the positively stained nuclei multiplied by the mean fluorescence-intensity value (MFI) of the positively stained nuclei.

#### Statistical Analysis

Data that were assumed to follow a log-normal distribution was log-transformed prior to statistical analysis. RT-qPCR was log-transformed and analyzed by Repeated Measurements Analysis of Variance (RM-ANOVA), or a mixed model if there was missing data, followed by Holm-Šídák’s multiple comparisons post-test. Scan^R data were log-transformed and analyzed with a paired t-test (two-sided). ELISA data was analyzed using a Wilcoxon matched-pairs signed-rank test. Data from WB were analyzed using 2-way ANOVA or mixed effect analysis. All graphs and analyses were generated with GraphPad Prism v10.1.2 (Dotmatics, Bishops Stortford, UK).

## Results

### P7-Pen inhibits expression and secretion of IRF5-dependent cytokines in human monocytes

We first evaluated the effect of the P7-Pen peptide on *IFNB* and proinflammatory cytokine mRNA expression in primary human monocytes isolated from buffy coats of healthy donors and stimulated with the TLR7 ligand R837 or several synthetic TLR7/8 or TLR8 ligands (Fig. 1A, Supplementary Fig. 1A). Monocytes were pre-treated with 15 μM P7-Pen or sterile endotoxin-free water as a solvent control. As a positive control for P7-Pen efficacy, cells were stimulated with ultrapure LPS, which confirmed significant inhibition of LPS-induced *IFNB*, *TNF*, and *IL1B* mRNA expression (Fig. 1A).

**Figure 1.**
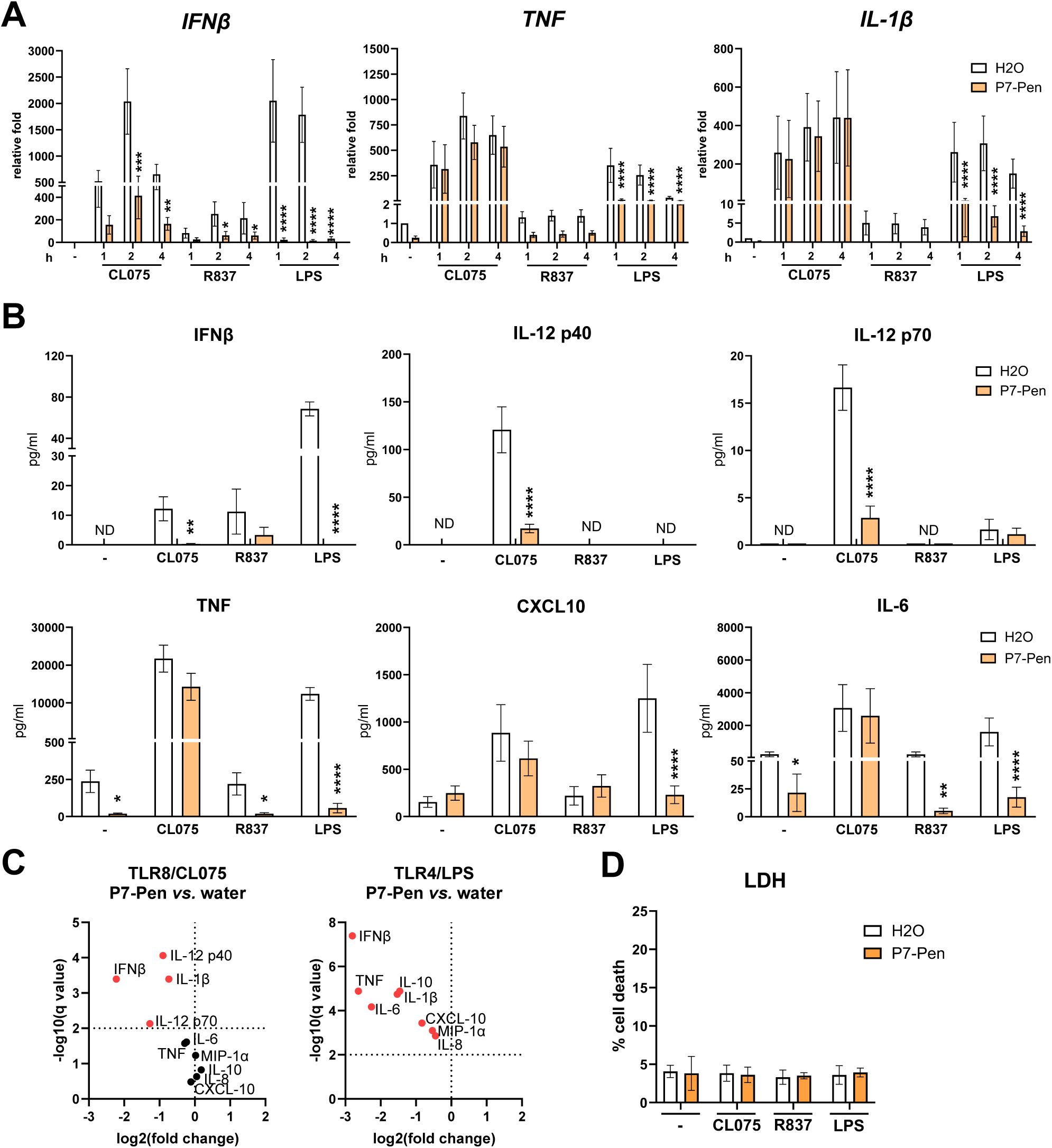
P7 inhibits TLR8--mediated IFNβ, IL-12p40 and IL-12 p70 secretion in human primary monocytes. Primary human monocytes isolated from PBMCs from healthy donors were pretreated with water solvent (H_2_O) or 15 μM P7-Pen for 30 min, followed by stimulation with CL075, R837 (1 μg/ml) or K12 LPS (100 ng/ml) for indicated time, followed by collection of supernatants and cell lysis. (**A**) Quantification of *IFNβ*, *TNF* and *IL-1β* mRNA expression by qRT-PCR in primary human monocytes. Data presented as relative fold change when compared to unstimulated sample pretreated by water (H_2_O), mean ± SEM (n = 6–11). (**B, C**) Cytokine secretion was addressed by ELISA for IFNβ and by BioPlex for other cytokines and graphed as mean ± SEM (n = 5–8). (**C**). P7-Pen effect on CL075- or LPS-mediated secretion of all tested cytokines shown as Volcano plots generated using multiple paired t-test analysis, controlled for a False Discovery Rate (FDR) of 1%. (**D**) Cell viability was addressed by LDH content in supernatants using LDH cytotoxicity assay and presented as % of dead cells, mean ± SD. (**A, B, D**) Statistical testing was done by two-way ANOVA or mixed effects model on log-transformed data (\**P* < 0.05, \*\**P* < 0.01, \*\*\**P* < 0.001, \*\*\*\**P* < 0.0001).

P7-Pen significantly inhibited R837- and CL075-induced *IFNB* expression but had no effect on CL075-mediated *TNF* or *IL-1B* mRNA levels (Fig. 1A). Similar results were observed with the TLR8 ligand TL8-506 and the dual TLR7/8 ligand R848 (Supplementary Fig. 1A). Consistent with mRNA data, P7-Pen significantly reduced LPS-induced secretion of IFNβ and proinflammatory cytokines (Fig. 1B, 1C). In contrast, P7-Pen selectively inhibited TLR8-induced IRF5-dependent cytokines—IFNβ, IL-12p40, and IL-12p70—and did not affect proinflammatory cytokine secretion (Fig. 1B, 1C, Supplementary Fig. 1B). LDH release assay did not indicate any cell cytotoxicity by the treatments (Fig. 1D).

As the levels of secreted IFNβ following TLR7 and TLR8 stimulation were in general relatively low (Fig. 1B), we investigated STAT1 (Y701) phosphorylation as an indirect readout to assess the peptide effects on IFNβ signaling. Western blot analysis showed that, in addition to LPS, all tested TLR7/8 ligands (R848, CL075, TL8-506, and R837) induced detectable STAT1 phosphorylation, which was consistently reduced by P7-Pen (Supplementary Fig. 1C, D).

While P7-Pen blocks TIRAP–MyD88 interaction, the small-molecule inhibitor TAK-242 (Resatorvid) specifically disrupts TLR4 interactions with TIRAP and TRAM (21). To confirm that P7-Pen’s inhibition of TLR8-mediated IFNβ expression is TIRAP-specific and not related to TLR4, we compared IFNβ and TNF mRNA expression in monocytes stimulated with LPS or CL075 and pre-treated with either TAK-242 or P7-Pen. Only P7-Pen, not TAK-242, significantly reduced CL075-induced IFNβ expression, while both inhibitors effectively blocked LPS-induced IFNβ and TNF expression (Supplementary Fig. 1E).

In summary, these results demonstrate that P7-Pen selectively inhibits TLR7- and TLR8-mediated IFNβ expression and secretion and TLR8-induced IL-12p40 and IL-12p70 production in primary human monocytes.

### P7-Pen inhibits TLR4- but not TLR8-mediated responses in human neutrophils

Neutrophils are the most abundant immune cells in human blood and serve as a first line of defense against bacterial infections (22). They are rapidly recruited from the bone marrow to infection sites, where they eliminate pathogens through phagocytosis, reactive oxygen species production, and NET formation. In addition to these effector functions, neutrophils express TLR2, TLR4, TLR6 and TLR8—and respond to LPS and synthetic TLR8 ligands with secretion of proinflammatory cytokines such as TNF, IL-6, and IL-8 (23, 24). Neutrophils are recruited to the inflammation site rapidly and in a large number, thus it was interesting to evaluate whether P7 affects neutrophils similar to monocytes.

To test whether P7-Pen modulates neutrophil cytokine responses, freshly isolated neutrophils from healthy donors were pre-treated with P7-Pen, solvent control (water), or TLR8 antagonist CU-CPT9a (5 μM), followed by stimulation with LPS or CL075 for 4 or 20 hours. Cytokine secretion was measured by ELISA (Supplementary Fig. 2A, B). Cell viability was assessed by LDH assay and showed no significant cytotoxicity (Supplementary Fig. 2C). As expected, CU-CPT9a efficiently inhibited CL075-induced TNF, IL-6, and IL-8 secretion at both time points, confirming TLR8 involvement. In contrast, P7-Pen did not affect CL075-mediated cytokine production (Supplementary Fig. 2A), while significantly suppressing LPS-induced secretion of TNF, IL-6, and IL-8 (Supplementary Fig. 2B), consistent with its inhibitory effect on TLR4 signaling previously observed in monocytes (Fig. 1B, C).

In summary, P7-Pen selectively inhibits TLR4-mediated cytokine responses in neutrophils but does not interfere with TLR8-driven proinflammatory signaling.

### P7-Pen inhibits TLR7- and TLR8-mediated IFNβ secretion in the human whole blood model

The whole blood model offers a physiologically relevant system to study TLR7- and TLR8-mediated responses due to the presence of all immune cell types, including dendritic cells— key producers of type I IFNs such as IFNβ in response to viral nucleic acids (25). Notably, IFNβ secretion in response to the TLR7/8 ligand CL075 was higher than that induced by LPS, likely reflecting the combined contribution of TLR7- and TLR8-expressing cells (e.g., monocytes, plasmacytoid, and conventional dendritic cells).

We employed a previously established human whole blood model using lepirudin as an anticoagulant to preserve complement activity (20). Given our earlier findings that TLR8 mediates IFNβ secretion in response to *S. aureus* in human cells, we tested whether P7-Pen could inhibit IFNβ production triggered by CL075, R837, or *S. aureus* particles (Fig. 2A). LPS and *E. coli* particles were included as controls to confirm peptide efficacy, as P7-Pen is known to inhibit cytokine responses mediated by these stimuli (16).

**Figure 2.**
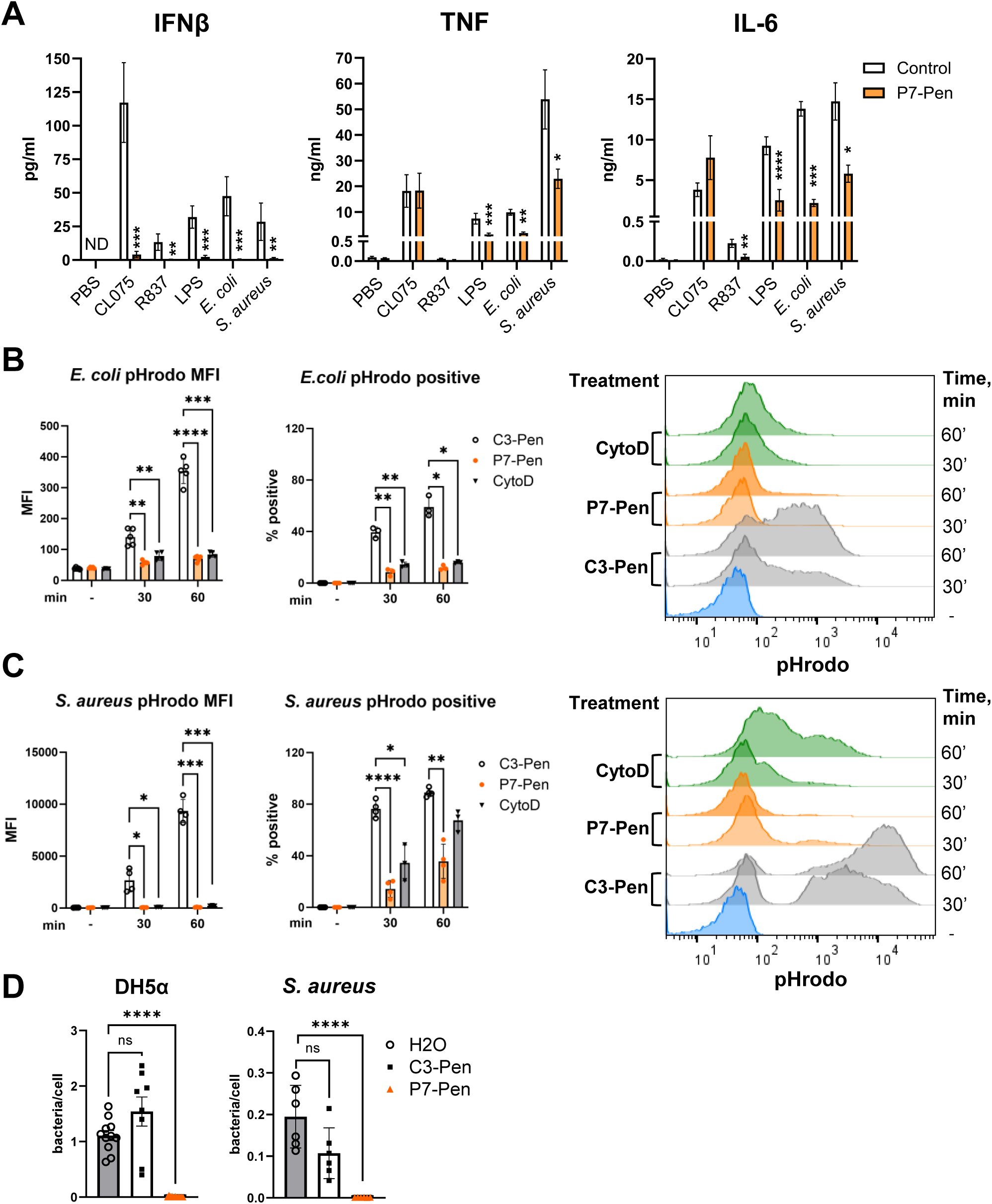
P7 inhibits IFNβ release downstream TLR7/8 in a human whole-blood model and decreases the uptake of *S. aureus* particles and live bacteria by human monocytes. (**A**) Whole-blood assay for samples of healthy donors with lepirudin as an anticoagulation reagent. Blood samples were pretreated with water solvent or 20 μM P7-Pen peptide for 30 min, followed by addition of CL075, R837 (2 μg/ml), K12 LPS (100 ng/ml), *E. coli* particles (2x10^6^/ml), or *S. aureus* particles (4x10^6^/ml) for 5 h before the collection of plasma samples. Plasma samples were probed for IFNβ, TNF and IL-6 secretion by ELISA. Data presented as mean ± SEM, statistical significance evaluated using Wilcoxon matched-pairs signed-rank test. (**B, C**) Quantification of phagocytosis based on flow cytometry for primary human monocytes pretreated by a solvent (water, H_2_O) or 15 μM P7-Pen, or 3 μM CytoD for 30 min and incubated with *E. coli* (**B**) or *S. aureus* (**C**) pHrodo particles for indicated time. Median pHrodo fluorescence intensity MFI and percentage of pHrodo-positive cells shown on graphs (**B, C**). Data presented as mean ± SEM, statistical significance evaluated by 2way ANOVA or mixed effect analysis. Representative images of flow cytometry results for one out of 3-5 donors are shown on the right. (**D**) Results of live bacteria phagocytosis assays for DH5α *E. coli* and *S. aureus* Wood strain by PMA-differentiated THP-1 monocytic cells. Data presented as mean ± SD, statistical significance evaluated by unpaired t-test (**A-D**) Significance levels: \**P* < 0.05, \*\**P* < 0.01, \*\*\**P* < 0.001, \*\*\*\**P* < 0.0001, nonsignificant if not shown otherwise; ND, not detected.

Consistent with previous results, P7-Pen significantly reduced IFNβ secretion in response to LPS and *E. coli*, and peptide in addition inhibited CL075-, R837-, and *S. aureus*-mediated IFNβ release (Fig. 2A). R837 did not induce notable proinflammatory cytokine secretion in this model, while CL075-mediated TNF and IL-6 production was unaffected by P7-Pen (Fig. 2A). Interestingly, P7-Pen significantly suppressed *S. aureus*-induced TNF and IL-6 secretion, despite having no effect on CL075-induced proinflammatory responses. IFNβ and, to some extent, TNF and IL-6 secretion in response to *S. aureus* are dependent on TLR8 (26), and, thus, requires efficient bacterial phagocytosis. We hypothesized that the inhibitory effect of P7-Pen may be due to the reduced bacterial uptake, leading to diminished delivery of bacterial ligands to endosomal compartments.

### P7-Pen inhibits the uptake of heat-killed and live S. aureus by human monocytes

We previously demonstrated that P7-Pen inhibits complement-independent uptake of *E. coli* by disrupting the TRAM–Rab11FIP2 interaction, a key step in recruiting Rab11FIP2 to bacterial particles (16). We have also shown that *Rab11FIP2* or *TRAM* silencing reduces the uptake of both *E. coli* and *S. aureus* by human macrophages (27).

Therefore, we compared the effect of P7-Pen on the uptake of *E. coli* (as a positive control) and *S. aureus* particles by primary human monocytes (Fig. 2B–D). Monocytes were pre-treated with 15 μM P7-Pen, scrambled control peptide (C3-Pen), or phagocytosis inhibitor Cytochalasin D (CytoD, 3 μM) before addition of fluorescent pHrodo-labeled bacterial particles. The pHrodo signal, detectable only in acidified phagosomes, was quantified by flow cytometry after 30 or 60 minutes of incubation (Fig. 2B, C). We found that P7-Pen significantly inhibited uptake of both *E. coli* and *S. aureus* particles, as shown by reduced median pHrodo fluorescence intensity (MFI) and decreased percentages of pHrodo-positive cells, comparable to CytoD treatment (Fig. 2C).

Since the uptake of bacterial particles and live bacteria may differ due to the active metabolic state, intact surface structures, and greater uptake efficiency of live bacteria (28), we also examined the effect of the peptide on live bacterial phagocytosis. In live bacterial uptake assays using PMA-differentiated THP-1 cells, pre-treatment with P7-Pen followed by co-incubation with live *E*. *coli* (DH5α) and *S*. *aureus* (protein A negative strain Wood 46) also led to a marked reduction in bacterial uptake compared to solvent (water) or control peptide treatments (Fig. 2D).

These findings support that the inhibition of *S. aureus*-induced cytokine secretion by P7-Pen in the whole blood model could be explained by impaired bacterial uptake.

### P7-Pen inhibits TIRAP recruitment to TLR8 Myddosome, IRF5 dimerization and nuclear translocation

We previously showed that TIRAP is recruited to the TLR8-Myddosome at a later stage, likely following the initial formation of the TLR8–MyD88–IRAK complex (14). This recruitment, observed around 45–60 minutes after ligand stimulation, coincides with a marked increase in CL075-induced posttranslational modification of IRAK1 (14). TIRAP silencing significantly reduces IRF5 nuclear translocation and the expression of IRF5-regulated genes such as *IFNB* and *IL12A* (14).

To test whether P7-Pen could block TIRAP recruitment to the TLR8-Myddosome, we performed immunoprecipitation (IP) assays using lysates from primary human monocytes (Fig. 3A). Endogenous TIRAP was immunoprecipitated using covalently bound anti-TIRAP antibodies from cells pre-treated with either P7-Pen or control peptide (C3-Pen) and stimulated with LPS (positive control) or CL075. In control-treated cells, TIRAP co-precipitated with MyD88, IRAK4, and IRAK1 following stimulation. At 60 minutes post-stimulation, we observed enhanced TIRAP recruitment to MyD88 and IRAK1. P7-Pen treatment strongly reduced TIRAP co-precipitation with both MyD88 and IRAK1, without affecting its interaction with IRAK4 (Fig. 3A).

**Figure 3.**
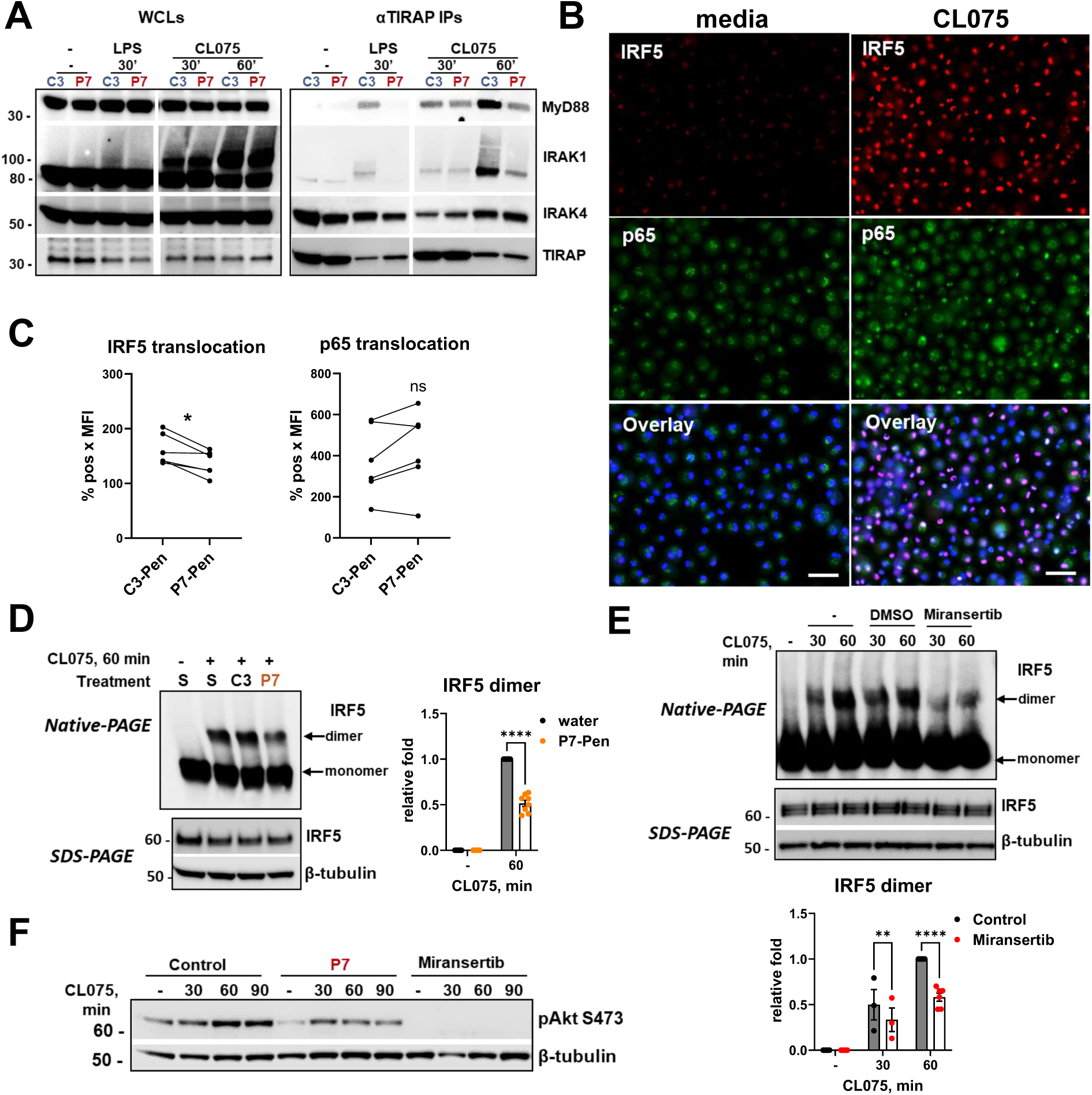
P7 interferes with TIRAP recruitment to TLR8-Myddosome, IRF5 dimerization and IRF5 nuclear translocation in human monocytes. (**A-C**) Primary human monocytes were pretreated with 15 μM C3-Pen (control) or P7-Pen for 30 min, followed by stimulation with CL075 (1 μg/ml) or LPS (100 ng/ml) for indicated time. (**A**) Endogenous TIRAP was immunoprecipitated from lysates (whole cell lysates – WCLs) of stimulated cells (500 μg protein/IP). LPS stimulation was applied as a positive control for TIRAP recruitment to the activated Myddosome. Cellular lysates were analyzed in parallel as input control (7.5%), with WB performed for total levels of MyD88, IRAK1, IRAK4, and TIRAP. A representative experiment from three consecutive experiments with different donors is shown, and subfigures represent different parts of the same WB membrane. (**B, C**) Monocytes were fixed and double stained for IRF5 and NF-kB (p65/RelA), DNA stained by Hoechst 3342 for nuclei visualization, and quantitative imaging performed by high-content screening (Olympus Scan^R system), and representative immunofluorescent images are shown (scale bar 50 µm). (**C**) The level of nuclear IRF5 and p65 was calculated as the percentage of positively stained nuclei multiplied by the mean fluorescence-intensity value (MFI) of the positively stained nuclei. In non-stimulated cells, the background-staining levels (%pos × MFI) for nuclear IRF5 and p65 were <15 and <73, respectively. (**D-E**) Western blot analysis of IRF5 using native and SDS-PAGE in lysates from human monocytes pretreated with control (water or DMSO solvents, control peptide C3-Pen), 15 μM P7-Pen peptide (**D, F**) or Akt inhibitor miransertib (2 μM) and stimulated by CL075 (1 μg/ml). The antibodies used are indicated on the figure, and β-tubulin probed as equal-loading control. (**D-E**) Representative images are shown for one of 3-6 donors. (**D, E**) Data on graphs for IRF5 dimer quantification presented as mean ± SEM, statistical significance evaluated by 2way ANOVA or mixed effect analysis. Significance levels: \*\**P* < 0.01, \*\*\*\**P* < 0.0001, nonsignificant if not shown otherwise. (**F**) Representative image for the samples used in (**D, E**) that shows the levels of pAkt S473 in control, P7-Pen or miranzertib treated monocytes.

Fluorescent staining and Scan^R high-throughput imaging analysis revealed that P7-Pen selectively inhibited IRF5, but not p65 NF-κB, nuclear translocation at 60 minutes following CL075 stimulation (Fig. 3B, 3C). This pattern closely mirrored the effects observed with *TIRAP* silencing (14).

IRF5 nuclear translocation is regulated by IKKβ, which phosphorylates IRF5 at Ser462— a key step required for IRF5 homodimerization and nuclear entry (7, 8). Unfortunately, specific commercial antibodies against IRF5 Ser462 (or Ser446 in isoform 1) phosphorylation are not currently available. Phos-tag gel analysis, which detects global IRF5 phosphorylation, did not show a clear reduction in overall IRF5 phosphorylation in either *TIRAP*-silenced (14) or P7-Pen-treated cells (Supplementary Fig. 3). Therefore, we assessed IRF5 homodimerization directly using native PAGE (Fig. 3D). P7-Pen pre-treatment significantly reduced IRF5 dimer formation compared to solvent (S) or control peptide (C3-Pen), correlating with reduced TIRAP recruitment (Fig. 3A) and IRF5 nuclear translocation (Fig. 3C).

Previously, we showed that TIRAP regulates IRF5 nuclear translocation in part via TIRAP-dependent Akt activation (14). Inhibition of Akt with specific inhibitors such as miransertib reduced both IRF5 nuclear translocation and IFNβ expression in monocytes (14). Consistently with these previous results, miransertib also significantly inhibited IRF5 dimerization (Fig. 3E). Similar to *TIRAP* silencing, P7-Pen markedly reduced Akt autophosphorylation at Ser473, and, as expected, this phosphorylation was fully abolished by miransertib (Fig. 3F). Interestingly, although P7-Pen only partially reduced Akt Ser473 phosphorylation, its effect on IRF5 dimerization was comparable to miransertib, suggesting a more complex mechanism underlying the peptide’s influence on IRF5 activation.

### P7-Pen inhibits TLR8-mediated Akt and IKKα/β phosphorylation in human monocytes

To examine how P7-Pen alters TLR8-driven signaling, we analyzed activation of key downstream molecules in monocytes stimulated with CL075 (Fig. 4A, B). In addition to suppressing Akt phosphorylation, P7-Pen—but not the control peptide C3-Pen—partially inhibited IRAK1 ubiquitination (visible as the upper ∼100 kDa band) and IKKα/β phosphorylation at Ser176/177, particularly at 60 minutes post-stimulation (and at 30 min and 60 min for phospho-IKKα/β). In contrast, TAK1 phosphorylation at Thr184/187 remained unaffected (Fig. 4A, B). Importantly, the Akt inhibitor miransertib did not affect IKKα/β phosphorylation (Fig. 4C), suggesting that P7-Pen’s inhibition of IKKα/β is not mediated through Akt, but likely results from reduced upstream IRAK1 activation. Interestingly, although P7-Pen impaired IRAK1 ubiquitination and IKKα/β phosphorylation, it did not alter early TAK1 phosphorylation, which peaked at 15 minutes (Fig. 4A, B). Because TIRAP accumulates at the Myddosome only at 45-60 min after TLR8 stimulation (14), early TAK1 activation may be TIRAP-independent.

**Figure 4.**
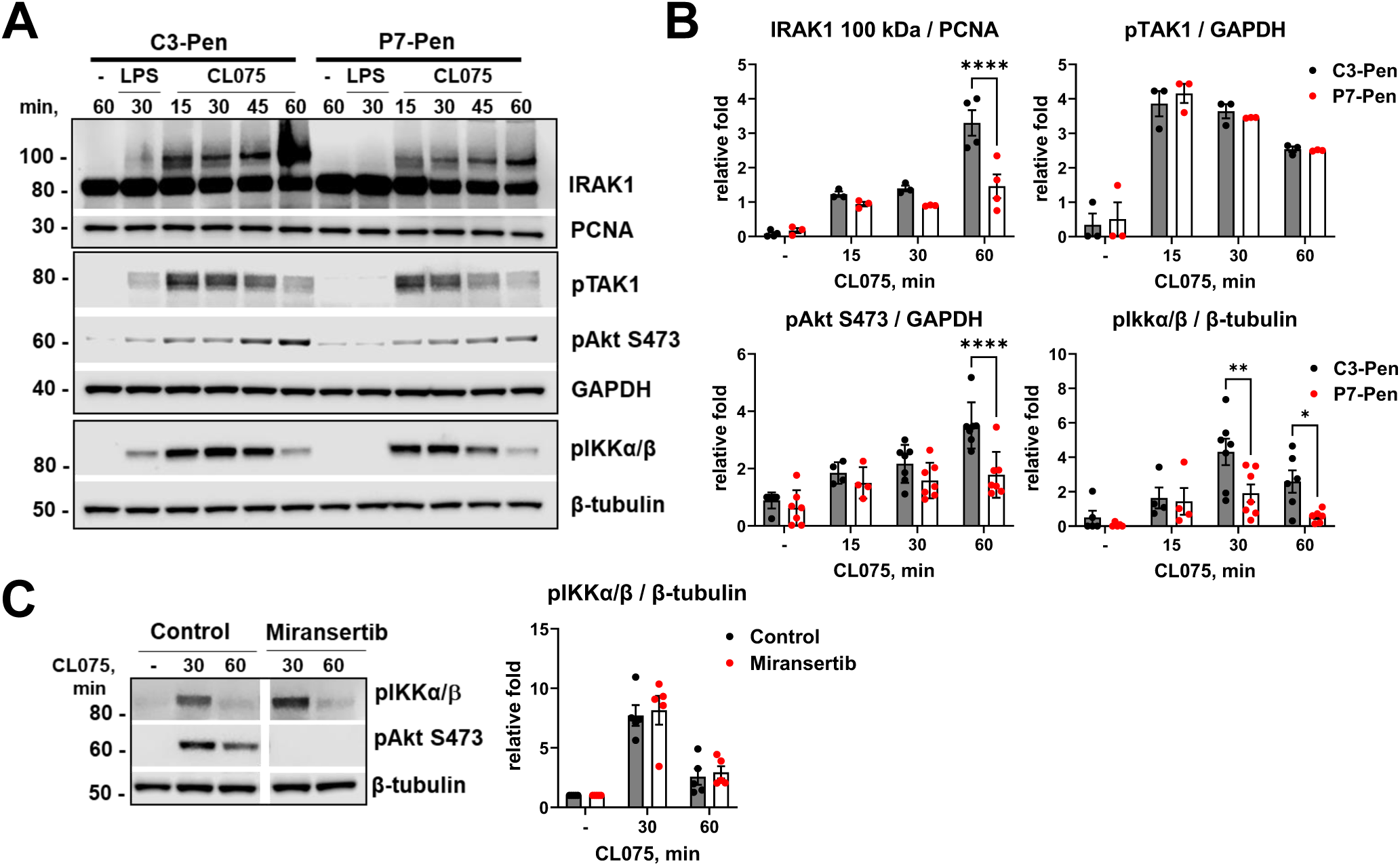
Pre-treatment of human monocytes by P7-Pen significantly reduced TLR8-mediated ubiquitination of IRAK1, and phosphorylation of Akt and IKKα/β. The antibodies used are indicated in the figure. PCNA, GAPDH or β-tubulin were used as equal-loading controls. (**A**) Primary human monocytes (n = 3-7 donors) were pretreated with 15 μM control peptide or P7-Pen for 30 min, followed by stimulation with CL075 (1 μg/ml) or LPS (100 ng/ml) for indicated time, followed by cell lysis and WB analysis. (**C**) Monocytes (n = 5 donors) were pretreated with DMSO (solvent control) or Akt inhibitor Miransertib (2 μM) and stimulated by CL075 (1 μg/ml) for indicated time, followed by cell lysis and WB analysis. (**B, C**) Graphs show quantification of protein levels relative to GAPDH, PCNA or β-tubulin. Data on graphs (**B, C**) presented as mean ± SEM, statistical significance evaluated by 2way ANOVA or mixed effect analysis. Significance levels: \**P* < 0.05, \*\**P* < 0.01, \*\*\*\**P* < 0.0001, nonsignificant if not shown otherwise.

### P7-Pen does not inhibit TLR7-mediated signaling in murine primary cells

In the current study, we demonstrated that P7-Pen inhibited IFNβ expression and secretion downstream of human TLR7 and TLR8 (Figs. 1, 2) and prevented TIRAP recruitment to the endosomal TLR8-Myddosome (Fig. 3A). Previous studies have shown that TIRAP knockout (KO) in murine bone marrow-derived macrophages (BMDMs) impairs murine TLR7- or TLR9-mediated *Ifnb* expression (12, 13). Direct TIRAP recruitment to murine TLR7 (mTLR7) in wild-type BMDMs has still not been examined. Given the relevance of murine models in studying TLR7-driven diseases, we thus tested whether P7-Pen could inhibit mTLR7-mediated responses.

P7-Pen effectively inhibited R848-induced IFNβ expression, secretion, and STAT1 phosphorylation in human monocytes (Supplementary Fig. 1A–D). Since R848 is an effective agonist for mTLR7 (29), it was used for stimulation of murine cells, with LPS serving as a positive control. Murine bone marrow (BM) cells were isolated from C57BL/6J mice and differentiated into either BMDMs (using M-CSF) or immature dendritic cells (iDCs, using GM-CSF + IL-4), as these subsets may differ in their endosomal TLR responsiveness (30). We also tested immortalized BMDMs (iBMDMs), which were used before to demonstrate the efficacy of TLR4 inhibition by P7-Pen peptide (16). Cells were pre-treated with P7-Pen or solvent control (water), then stimulated with R848 (Fig. 5A–C) or LPS (Supplementary Fig. 4). While P7-Pen efficiently inhibited LPS-induced expression of *Ifnb*, *Tnf*, and *Il-1b* (Supplementary Fig. 4), it had no inhibitory effect on R848-induced cytokine expression in primary BMDMs, iDCs or iBMDMs (Fig. 5A–C). A reduction in *Il-1b* expression was observed in P7-Pen-treated BMDMs, but this effect was not replicated in iDCs or iBMDMs (Fig. 5A–C).

**Figure 5.**
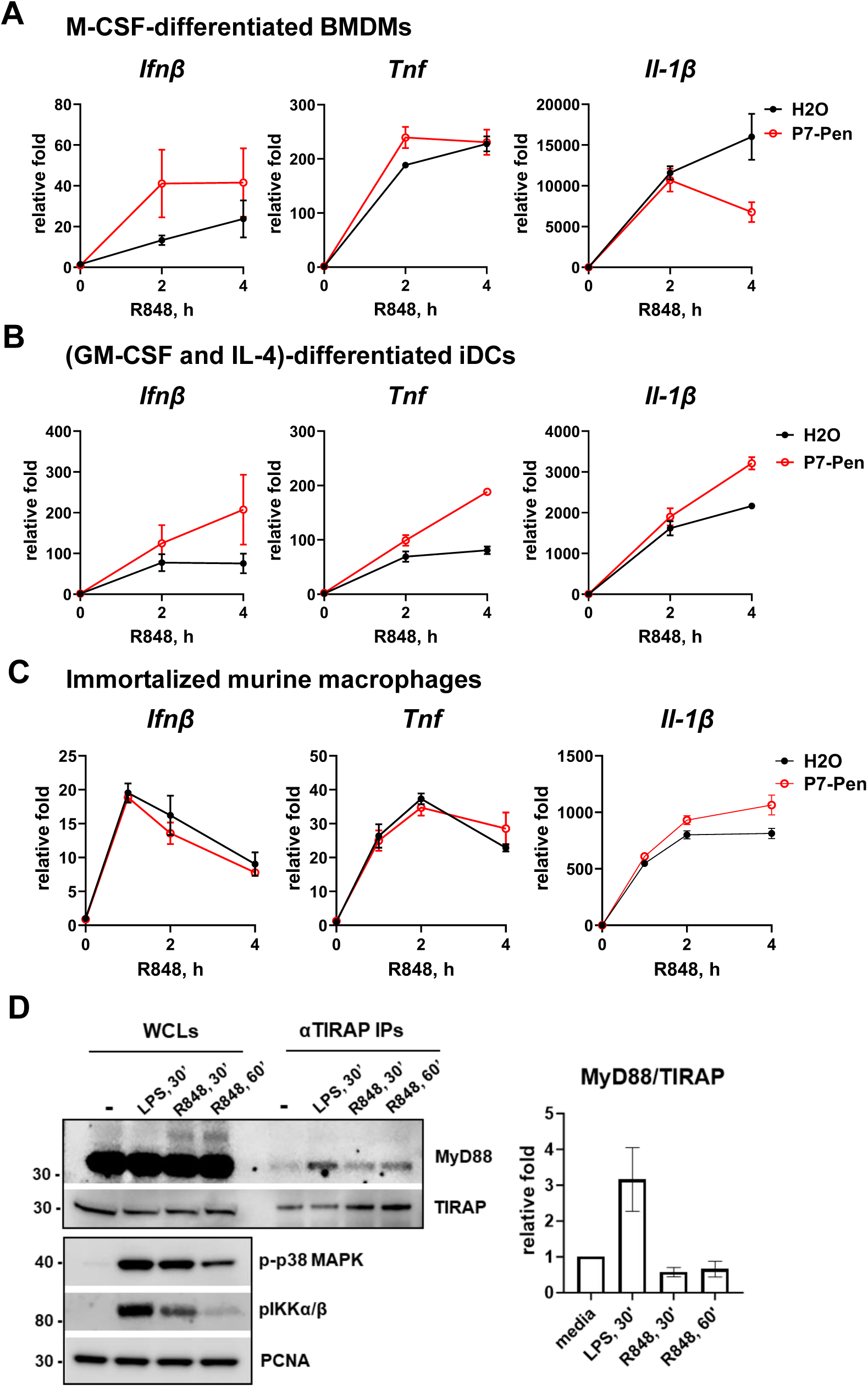
P7 peptide does not inhibit TLR7-mediated cytokine expression and TIRAP is not recruited to TLR7-Myddosome in murine macrophages. BMDMs (**A**) and iDCs (**B**) were generated from BM cells from C57BL/6 mice. (**A-C**) M-CSF-differentiated BMDMs (**A**), bone marrow derived iDCs (GM-CSF and IL-4-differentiated) (**B**) or immortalized murine macrophages (**C**) were pretreated with water solvent (H_2_O) or 10 μM P7-Pen for 30 min and stimulated with R848 (1 μg/ml) for indicated time, followed by cell lysis, RNA isolation and RT-qPCR analysis of *Ifnβ, Tnf* and *Il-1β* mRNA expression. Data presented as relative fold change when compared to unstimulated sample pretreated by water (H_2_O), mean ± SEM (n = 3). (**D**) Immortalized murine macrophages were kept untreated or stimulated with B4 LPS (100 ng/ml) or R848 (1 μg/ml) for indicated time, followed by cell lysis and IP assays. Endogenous TIRAP was immunoprecipitated from WCL of stimulated cells (1 mg protein/IP). LPS stimulation was applied as a positive control for TIRAP recruitment to MyD88. Cellular lysates were analyzed in parallel as input control (5%), with WB performed for total levels of MyD88 and TIRAP in WCLs and IPs, and for levels of phosphorylated p38 MAPK and IKKα/β in WCLs as a positive control for cell activation by TLR ligands. A representative experiment is shown from a total of three consecutive experiments.

We next investigated whether TIRAP is recruited to the mTLR7–Myddosome by performing endogenous TIRAP IPs from lysates of immortalized murine macrophages stimulated with LPS (positive control) or R848 (Fig. 5D). While both LPS and R848 stimulation induced phosphorylation of IKKα/β and p38 MAPK, only LPS triggered enhanced TIRAP recruitment to MyD88 (Fig. 5D). This lack of TIRAP recruitment may explain why P7-Pen fails to inhibit R848-induced *Ifnb* expression in murine cells.

## Discussion

We have previously demonstrated that TIRAP is required for optimal expression and secretion of IRF5-dependent cytokines downstream human TLR8, which is mediated by the recruitment of TIRAP to the TLR8–MyD88 complex (14). In primary human cells, TIRAP is recruited to the TLR8-MyD88-IRAKs (TLR8-Myddosome) approximately one hour after TLR8 activation, leading to increased co-precipitation of IRAK1 and MyD88 with TIRAP, and activation of the Akt kinase (14). We hypothesize that TIRAP may facilitate IRAK1 activation by IRAK4 during the later stages of TLR8-Myddosome formation, or mediate stabilization and prolongation of signaling from this signaling complex. Through its regulatory role in TLR8-Myddosome dynamics and the induction of Akt kinase activity, TIRAP recruitment contributes to enhanced IRF5 activation, nuclear translocation, and the expression of IRF5-regulated genes (14).

Here, we demonstrate that the P7-Pen peptide, which interferes with the TIRAP–MyD88 interaction (16), inhibits the recruitment of TIRAP to the TLR8-Myddosome and reduces TLR8-mediated IFNβ expression and secretion in human monocytes. Notably, the small molecule inhibitor TAK-242, which blocks the TLR4–TIRAP interaction and TLR4-mediated signaling (21), had no effect on TLR8-induced cytokine expression. However, reducing the TIRAP–MyD88 interaction with P7-Pen inhibited both TLR4-induced cytokine expression and TLR8-mediated IFNβ expression. In this study, we tested and compared several TLR7/8 ligands (CL075, R848), as well as TLR7-specific (R837) and TLR8-specific (TL8-506) ligands and observed consistent inhibition of IFNβ expression and secretion by P7-Pen across all tested stimuli. Consistent with previous findings on TIRAP silencing, P7-Pen also inhibited TLR8-mediated secretion of IL-12p40 and IL-12p70 by human monocytes.

While our previous study focused on the role of TIRAP in TLR8 signaling in monocytes-derived macrophages (MDMs), we did not examine its potential involvement in TLR7-driven responses (14) since MDMs have low levels of TLR7, and are practically non-responsive to R837 (31). Therefore, we could not test the impact of *TIRAP* knockdown on TLR7-mediated responses in MDMs. In this study we instead examined the responses of primary monocytes using the inhibitory peptide. Monocytes exhibited weak responses to the TLR7 ligand R837 compared to the TLR8-specific ligand TL8-506, with almost no induction of proinflammatory cytokines expression following R837 stimulation. Nevertheless, R837-induced IFNβ expression and secretion was detectable, as was also STAT1 phosphorylation, indicative of type I IFN receptor signaling. In the whole blood model, the response to R837—measured by IFNβ secretion—was more robust, likely due to the plasmacytoid dendritic cells expressing higher levels of functional TLR7 (32, 33). Our data clearly demonstrated that the P7-Pen peptide inhibits both TLR7- and TLR8-mediated IFNβ expression and secretion in human monocytes and peripheral blood. The similar inhibitory effect of P7-Pen on TLR7- and TLR8-mediated IFNβ expression is indicative of a shared mechanism of IFNβ regulation by these two TLRs.

We previously showed that TLR8 is a key sensor of Gram-positive bacteria such as *S. aureus* in human monocytes and MDMs (9, 26, 34), and contributes to IL-8 release by neutrophils during phagocytosis (35). Expression of *IL-12A* and *IFNβ* in response to *S. aureus* is highly TLR8 dependent (34), and *TIRAP* silencing reduces *IFNβ* secretion by infected macrophages (14). We therefore tested whether P7-Pen modulates *S. aureus*-induced cytokine production in whole blood, which reflects the combined response of TLR8-expressing monocytes, dendritic cells, and neutrophils (5). In this system, P7-Pen inhibited CL075- and R837-induced IFNβ secretion, without affecting TNF or IL-6 levels. In contrast, stimulation with *S. aureus* led to reduced secretion of IFNβ, TNF, and IL-6. Since TLR2 also contributes to TNF and IL-6 production in response to *S. aureus* (36), and TIRAP functions as a bridging adaptor for TLR2 (37), we considered whether this pathway might explain the broader effect. However, P7-Pen does not disrupt TLR2 signaling in monocytes and macrophages (16), suggesting another mechanism. We previously showed that P7-Pen inhibits complement-independent phagocytosis of Gram-negative bacteria by disrupting the TRAM–Rab11FIP2 interaction (16), and that Rab11FIP2 is required for phagocytosis of both Gram-negative and Gram-positive bacteria (27). Consistently, P7-Pen impaired the uptake of both *E. coli* and *S. aureus* by human monocytes. These findings suggest that reduced cytokine release upon *S. aureus* stimulation is likely mediated by impaired bacterial uptake, limiting RNA delivery to endosomal compartments and thereby dampening TLR activation.

Next, we investigated at which level the P7-Pen peptide interferes with molecular events initiated downstream of activated TLR8 in human monocytes. To ensure comparable signaling kinetics, we used the potent TLR7/8 ligand CL075, as previously applied in our study on *TIRAP* silencing (14). First, we provided direct evidence that P7-Pen inhibits TIRAP recruitment to the TLR8-Myddosome, which in turn leads to reduced late-stage IRAK1 ubiquitination and decreased phosphorylation of downstream IKKα/β at Ser176/177. TAK1 activation was not affected by the peptide, as full phosphorylation was already evident at 15 minutes post-stimulation and did not increase further during later stages associated with TIRAP recruitment. Overall, our findings suggest that the early-formed TLR8-Myddosome, lacking TIRAP, is sufficient for effective activation of downstream MAP kinases and NF-κB pathways required for proinflammatory cytokine expression. However, the activation and nuclear translocation of IRF5 may follow a distinct time kinetics and could involve additional upstream kinases beyond TAK1 for IKKα/β (Ser176/177) phosphorylation.

Both IKKα and IKKβ are required for the activity of the IKK complex (38, 39). While the catalytic activity of IKKα is not essential on its own, it has been suggested that, when associated with IKKβ, it may be activated by the latter and contribute to overall IKK complex activity (39, 40). Phosphorylation of IKKβ—but not IKKα—at Ser177 plays a central role in IKK complex activation by proinflammatory cytokines. This phosphorylation is mediated primarily by MAPK kinases such as TAK1 and, to a lesser extent, by trans-autophosphorylation (40).

Two independent research groups have demonstrated that IKKβ is the kinase responsible for phosphorylating IRF5 at Ser462/Ser446 (in IRF5 isoforms 2/1) downstream of TLR8 activation, a pre-requisite for IRF5 dimerization and nuclear translocation (7, 8). Due to the absence of site-specific antibodies, we were unable to directly assess phosphorylation at Ser462, and phos-tag gel analysis of total phospho-IRF5 provided limited information. However, IRF5 homodimerization (required for nuclear translocation) is strictly dependent on IKKβ-mediated phosphorylation (7). Thus, IRF5 homodimerization was evaluated as a functional readout of IRF5 activation, and shown to be significantly reduced by both P7-Pen and the Akt inhibitor.

We have previously demonstrated that TIRAP recruitment facilitates Akt activation and autophosphorylation at Ser473 to enhance IRF5 nuclear translocation (14). However, the precise mechanisms underlying Akt activation downstream TLR8 remains incompletely understood. Notably, PI3K/Akt activation downstream of IL-1β and TLR4 has been shown to depend on IRAK1 (41), raising the possibility that a similar mechanism may operate in the context of TLR8 signaling. Therefore, we suggest that TIRAP recruitment to the TLR8-Myddosome may promote Akt activation by enhancing IRAK1 activation.

Interestingly, treatment with the Akt inhibitor led to a marked reduction in IRF5 dimerization, a prerequisite for nuclear translocation, while leaving IKKα/β (Ser176/177) phosphorylation unaffected. Akt has been shown to phosphorylate IKKα at Thr23 downstream of TNF receptor signaling, a modification that positively regulates IKK complex activity (42, 43). We hypothesize that Akt may similarly regulate TLR8-induced IKK complex activity through phosphorylation of IKKα at Thr23, potentially enhancing the activity or stability of the IKKα/β heterodimer. However, due to the lack of functional commercial antibodies against this specific phospho-site, we were unable to directly assess the effects of either the Akt inhibitor or the P7-Pen peptide on IKKα Thr23 phosphorylation downstream of TLR8 stimulation.

Overall, our data demonstrate that the P7-Pen peptide, by blocking TIRAP recruitment to the TLR8-Myddosome, inhibits late stage IRAK1 and Akt activation in response to TLR8 stimulation. This dual inhibition may contribute to the observed decrease in IKKα/β activity and subsequent impairment of IRF5 nuclear translocation. A schematic overview of the proposed model for TIRAP-dependent regulation of TLR8-mediated signaling—and the point of intervention by the peptide—is presented in Figure 6. We also propose a similar mechanism of TIRAP-mediated regulation of human TLR7-induced IFNβ expression.

**Figure 6.**
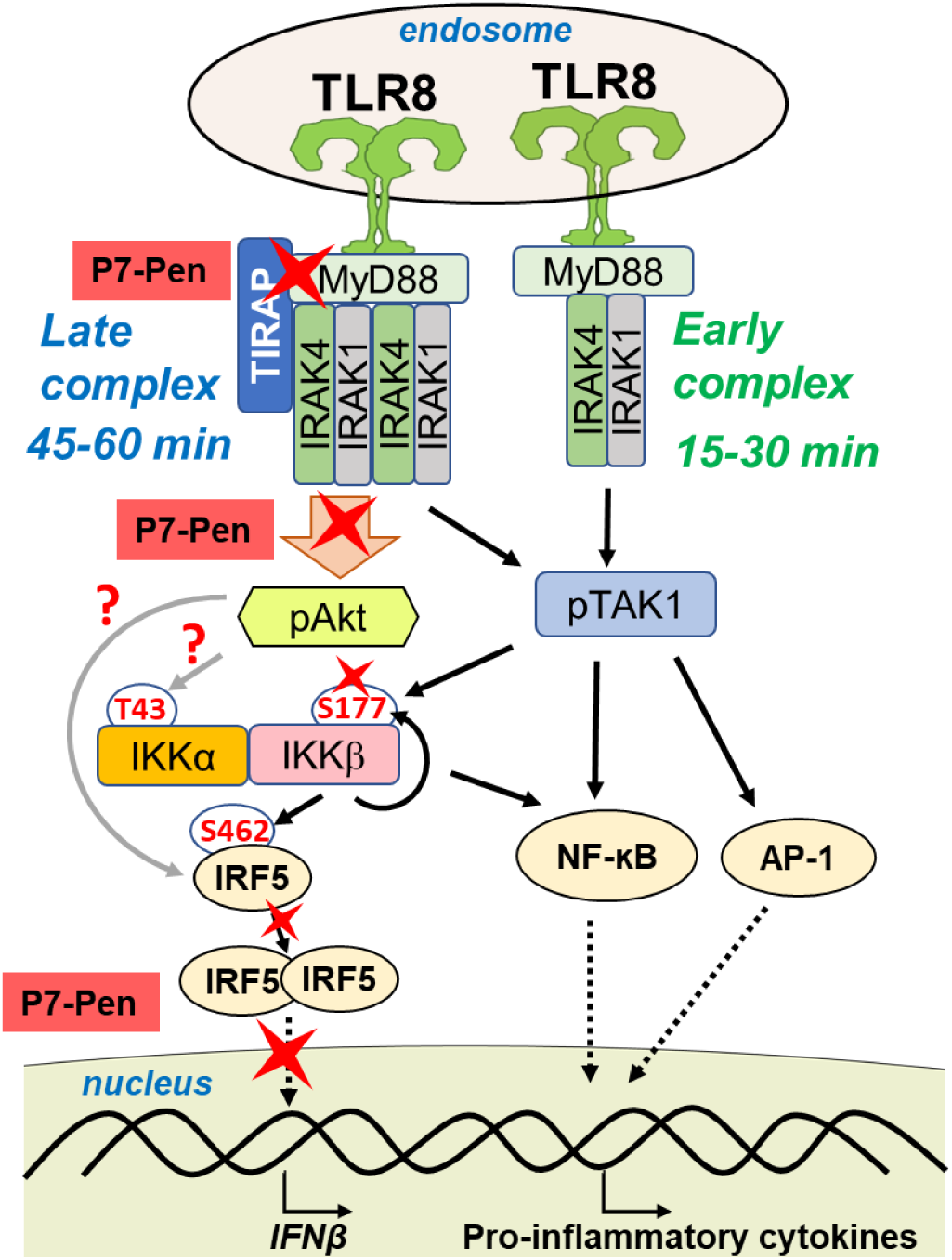
Mechanistic model for P7-Pen interacting partners and effects in TLR8-signaling. Recruitment of TIRAP to the TLR8-Myddosome complex 45–60 minutes after TLR8 activation (late complex) promotes Akt activation, IRF5 dimerization and nuclear translocation, and the expression and secretion of IFNβ and IL-12. The P7-Pen peptide disrupts TIRAP recruitment to the TLR8-MyD88 complex, thereby inhibiting phosphorylation of Akt and IKKα/β, IRF5 dimerization, and nuclear translocation, ultimately leading to reduced expression of IRF5-regulated genes.

Interestingly, P7-Pen failed to inhibit, and even enhanced *Ifnb* expression downstream of murine TLR7 in primary murine cells. Moreover, stimulation of murine iBMDMs with the TLR7 ligand R848 did not lead to increased recruitment of TIRAP to MyD88, whereas LPS did. This was intriguing considering a previous report showing that iBMDMs from TIRAP knockout (KO) mice had reduced *Ifnb* induction upon R848 stimulation (13). However, the direct recruitment of TIRAP to murine TLR7 was not previously addressed, and the results obtained using TIRAP KO cells may reflect broader changes in cellular signaling properties due to compensatory adjustments following the genetic deletion, such as altered expression of other signaling molecules. To definitively determine the role of TIRAP in regulating murine TLR7 signaling, it would be valuable to directly compare responses in wild-type, TIRAP KO and silenced cells, and address TIRAP localization following TLR7 activation using confocal microscopy. Based on our current findings, we propose that TLR7 signaling in murine macrophages differs from human cells and is potentially independent of TIRAP recruitment.

Species-specific differences may further question the relevance of murine models for studies targeting TIRAP-regulated type I IFN secretion. Endosomal TLRs—such as TLR7, TLR8, and TLR9—have been shown to play a critical role in the development of the autoimmune disease systemic lupus erythematosus (SLE), and targeting these receptors, along with TLR-mediated type I IFN production, is considered a promising therapeutic strategy (44). Elevated TLR7 expression, independent of gene copy number, has been associated with more severe disease in human SLE patients, and a mutation in human TLR8 has been linked to severe autoimmune manifestations (45, 46). Notably, humanized models of SLE have also been established. These include the transfer of PBMCs from SLE patients into immunodeficient mice, or the transplantation of human hematopoietic stem cells to immunodeficient mice followed by intraperitoneal injection of pristane to induce lupus (reviewed in (47)). Such models may offer a more relevant platform for evaluating the efficacy of compounds that target human cell-specific regulatory pathways.

In addition to their roles in autoimmune disease pathogenesis (2, 30, 44, 48), TLR7 and TLR8 are also critically involved in immune responses to both viral and bacterial infections (9, 26, 34, 49–54). Cell type-specific differences in the expression levels of TIRAP and its utilization in TLR7 and TLR8 signaling, along with the potential to modulate type I IFN secretion using the novel SLAMF1-derived peptide, should be explored in more detail. Targeting TIRAP may hold therapeutic promise for a range of conditions, given the association of TIRAP single nucleotide polymorphisms with the incidence and severity of diseases such as tuberculosis, HIV, and systemic lupus erythematosus (SLE) (55), as well as other infections and inflammatory disorders in which TLR8 likely plays a role (56–58).Thus, understanding the contribution of TIRAP to the regulation of endosomal TLRs signaling, especially in human

immune cells, and finding tools/molecules to modulate this signaling might be of significant clinical relevance.

## Acknowledgements

This research was funded by the Research Council of Norway through its Centers of Excellence Funding Scheme, Grant 223255/F50 (to T Espevik), NTNU Discovery Grant 2020 (to T Espevik and M Yurchenko), the Liaison Committee for Education, Research and Innovation in Central Norway Innovation Researcher Grant 90794301 (to M Yurchenko) and Felles Forskningsutvalg (FFU) Grant 2022/2758 (to M Yurchenko).

## Author contribution

KEN: formal analysis, investigation, writing - editing; JS: methodology, resources, writing - editing; AS, IBM, SSB, MSG, CSG, KE, LR: investigation; TE: methodology, resources, writing - editing; MY - conceptualization, resources, formal analysis, supervision, funding acquisition, investigation, methodology, project administration, and writing - original draft, review, and editing.

