## Supplemental data for "A novel TIRAP-MyD88 inhibitor blocks TLR7 and TLR8-induced type I IFN responses"

### Supplementary data

#### SUPPLEMENTARY METHODS

##### *Reagents*

The TLR8 antagonist CU-CPT9a is previously described {Zhang, 2018 #93} and was provided by The Regents of the University of Colorado, a body corporate for and on behalf of the University of Colorado Boulder. TAK-242 (Resatorvid) selective TLR4 inhibitor was from MedChemExpress (#HY-11109).

##### *Phos-tag gel*

For total IRF5 phosphorylation analysis, SuperSep Phos-tag gels (50 mol/L, 7.5%, 17 well, 100 × 100 × 6.6 mm) (FUJIFILM Wako Pure Chemical Corporation, Osaka, Japan) were used with Running Buffer (1× Tris-Glycine with 0.1% SDS). Before transfer, gels were washed three times for with gel running buffer containing 20% methanol and 10 mM EDTA for 20 min and once with the same buffer without EDTA to facilitate the transfer procedure (according to manufacturer recommendations). GAPDH WB on normal SDS-PAGE gel was used for equal-loading control.

##### *Neutrophils isolation and TLR8 inhibitor for supplementary methods*

Human polymorphonuclear neutrophils were obtained from fresh peripheral blood drawn by venipuncture as described {Pouliot, 2002 #130}. Briefly, venous blood from healthy volunteers was collected in isocitrate anticoagulant solution and centrifuged at 250 × g for 10 min. Whole blood cells were obtained from the pellet following red blood cells (RBC) sedimentation in 2.0% Dextran T-500 (Sigma–Aldrich). Neutrophils were then separated by centrifugation on a 10 ml cushion of Lymphoprep™ (Axis-Shield). Contaminating RBC were removed by 20 seconds of hypotonic lysis in water, before adding concentrated HBSS (Sigma, Merck). Viability was higher than 98%, based on trypan blue dye exclusion and Cell Hostess Counter quantification (Thermo Fisher Scientific, Waltham, MA, USA). The entire procedure was carried out at RT under sterile conditions. Neutrophils were resuspended in RPMI with 10% human serum, and 5 × $10^6$ cells were seeded per well in 24-well culture plates (Costar). Some cells were pretreated with 15 μM P7-Pen peptide or TLR8 antagonist CU-CPT9a (5 μM) for 30 min before stimulation by TLR ligands. Cell free supernatants were collected after brief centrifugation of plates and stored at −80°C until analyses.

#### SUPPLEMENTARY FIGURES

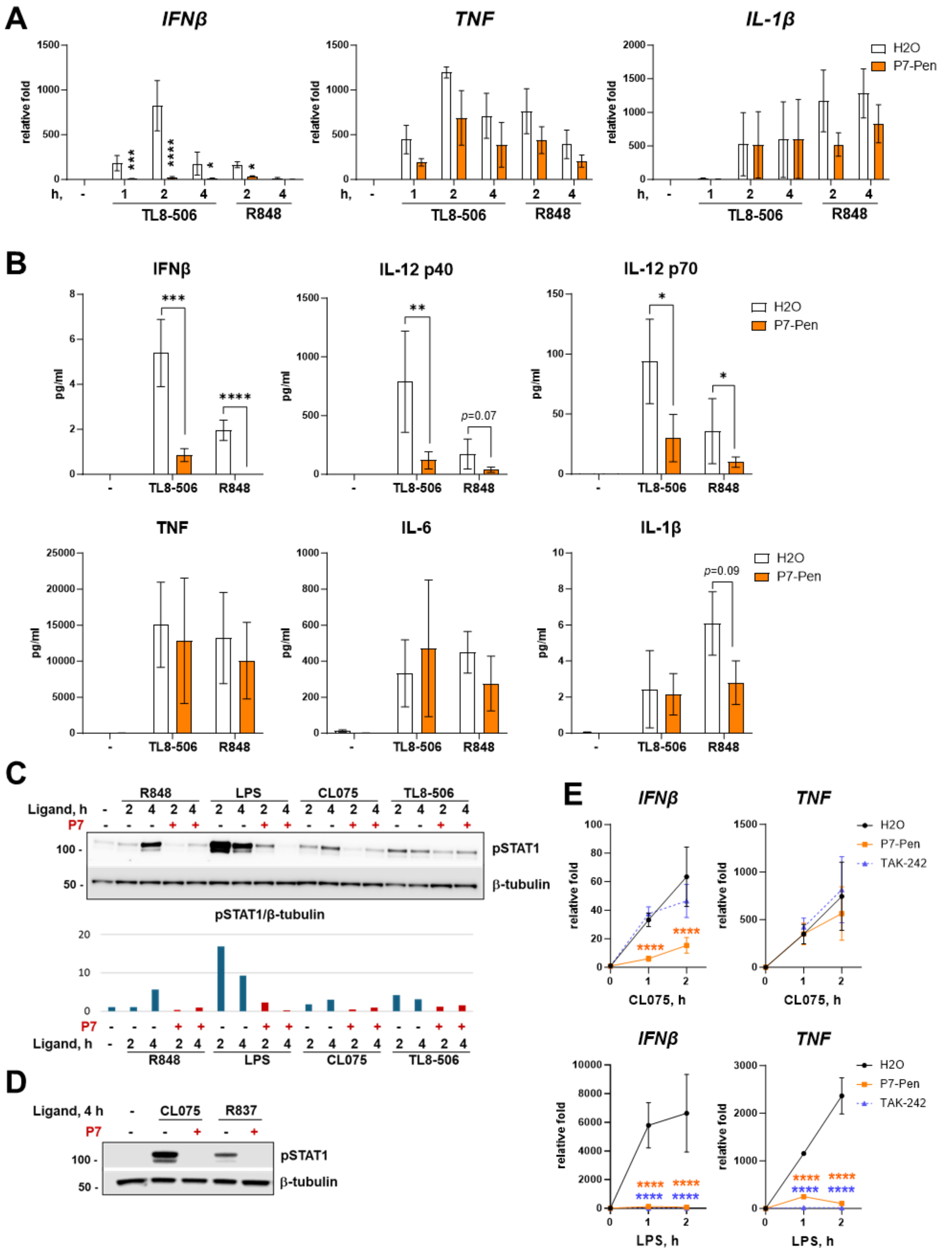


**Supplementary Figure 1. P7 inhibits IFNβ, IL-12p40, and IL-12 p70 secretion meditated by TL8-506/TLR8 and R848/TLR7/8 ligands in human monocytes.** Primary human monocytes isolated from PBMCs from healthy donors (n = 3) were pretreated with water solvent ($H_2O$) or 15 μM P7-Pen for 30 min, followed by stimulation with TL8-506 or R848 (1 μg/ml). (**A**) Quantification of *IFNβ*, *TNF* and *IL-1β* mRNA expression by qRT-PCR, presented as relative fold change when compared to unstimulated sample pretreated by water ($H_2O$), mean ± SEM. (**B**) Cytokine secretion addressed by ELISA for IFNβ and by BioPlex assays for IL-12 p40, IL-12 p70, TNF, CXCL10, IL-6 (mean ± SEM). (**C, D**) WB for pSTAT1 (Tyr701) levels in protein samples isolated along with RNA from Qiazol lysates (qPCR on Fig. 1A and Suppl. Fig. 1A) after stimulation of monocytes by different TLR ligands for 2 and 4 h. Graph in (**C**) shows pSTAT level normalized to endogenous control β-tubulin (quantification for representative image is shown, one out of three donors). (**E**) Monocytes (n = 3 donors) were pretreated with water solvent ($H_2O$), or 15 μM P7-Pen, or 10 μM TAK-242 for 30 min, stimulated by CL075 (1 μg/ml) or LPS (100 ng/ml) for 1 or 2 h, followed by cell lysis, RNA isolation and quantification of *IFNβ* and *TNF* expression by RT-qPCR. Data presented as relative fold change when compared to unstimulated sample pretreated by water ($H_2O$), mean ± SEM. (**A, B, E**) Statistical testing was done by two-way ANOVA on log-transformed data (*$P < 0.05$, **$P < 0.01$, ***$P < 0.001$, ****$P < 0.0001$, and otherwise non-significant).

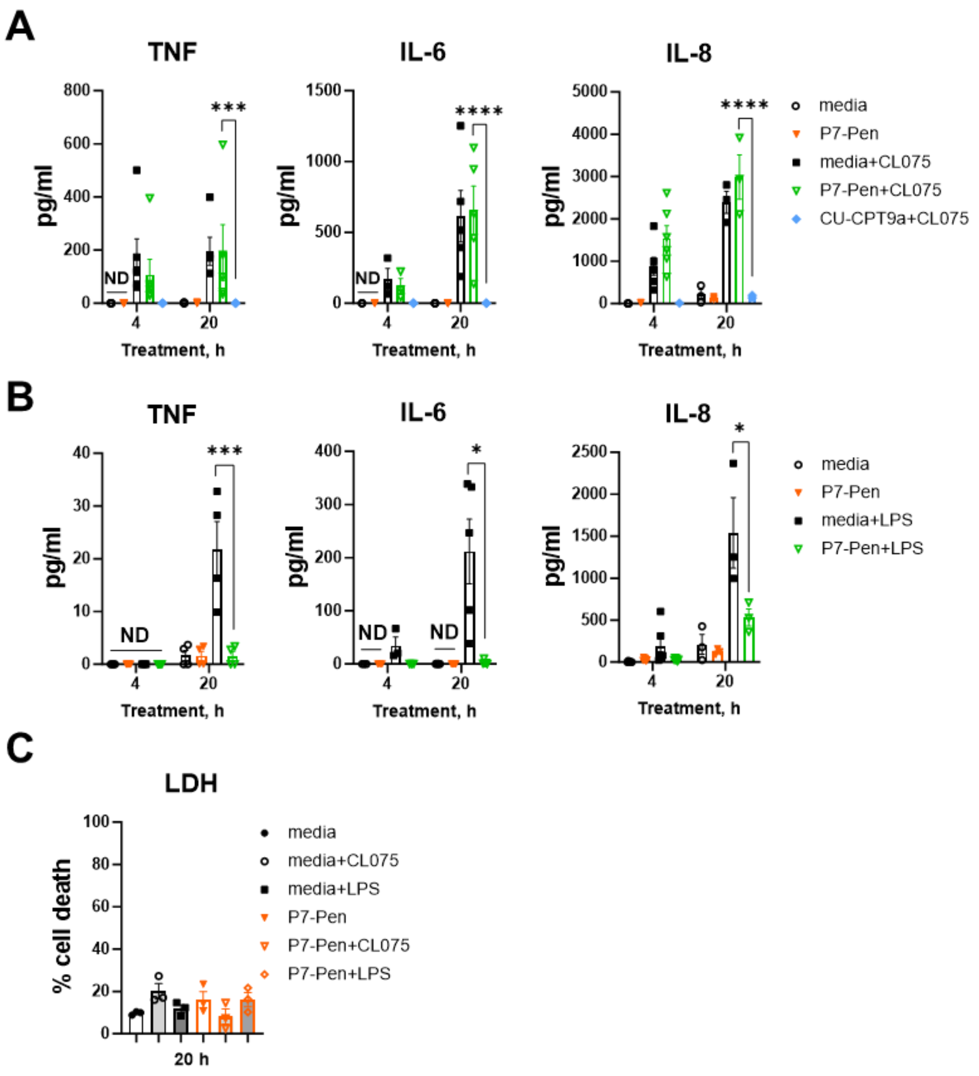


**Supplementary Figure 2. P7 inhibits TLR4/LPS but not TLR8/CL075-mediated pro-inflammatory signaling in human neutrophils.** Primary human neutrophils were freshly isolated from blood samples of healthy donors (n = 3-6) and immediately used for experimental procedures. Cells were seeded in 24-well plates, $5 \times 10^6$ cells/well, pretreated with 15 µM P7-Pen or CU-CPT9a TLR8 antagonist (5 µM) for 30 min, followed by stimulation with CL075 (1 µg/ml) or LPS (100 ng/ml) for 4 or 20 h. At the end of stimulation time, plates were placed, and supernatants collected for ELISA or LDH assay. (**A, B**) Cytokine secretion of TNF, IL-6 or IL-8 mediated CL075 (**A**) or LPS (**B**) were addressed by ELISA and graphed as mean ± SEM, with individual values indicated. Statistical testing was done by two-way ANOVA on log-transformed data, significance levels: *$P <$ 0.05, ***$P <$ 0.001, ****$P <$ 0.0001, nonsignificant if not shown otherwise; ND, not detected. (**C**) Cell viability was addressed by LDH content in the selected supernatants (n = 3 donors) using LDH cytotoxicity assay and presented as % of dead cells, mean ± SD.

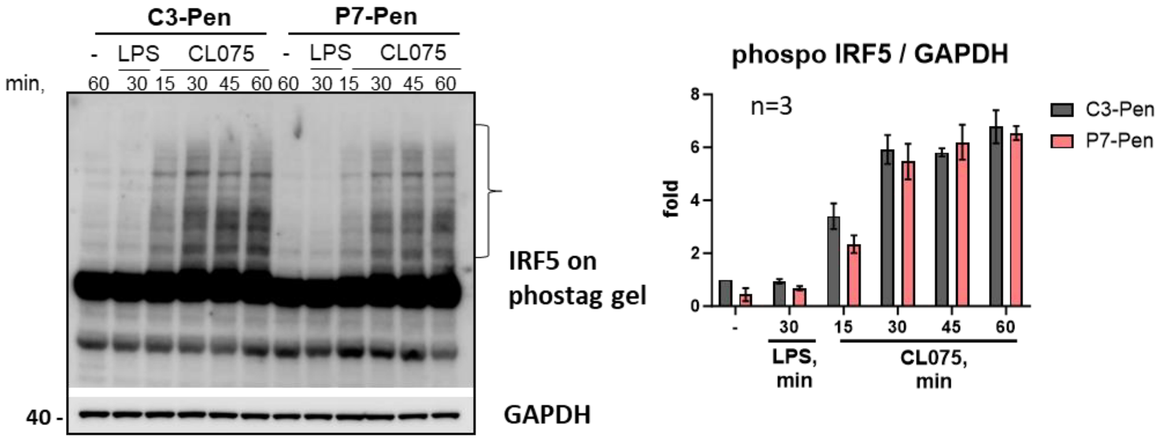


**Supplementary Figure 3. Total phosphorylation pattern of IRF5 in CL075-stimulated cells is unaltered by P7-Pen treatment.** Lysates were resolved using PhosTag gel as recommended by manufacturer, followed by Western blot analysis using total IRF5 Abs. GAPDH Western blot (lower panel) in parallel conventional SDS-PAGE was used for loading control. Quantification of total phospho-IRF5 bands normalized to GAPDH loading control for three independent experiments (n = 3 donors) is shown as a bar plot and did not reveal significant differences for C3-Pen and P7-pen treated cells (ANOVA).

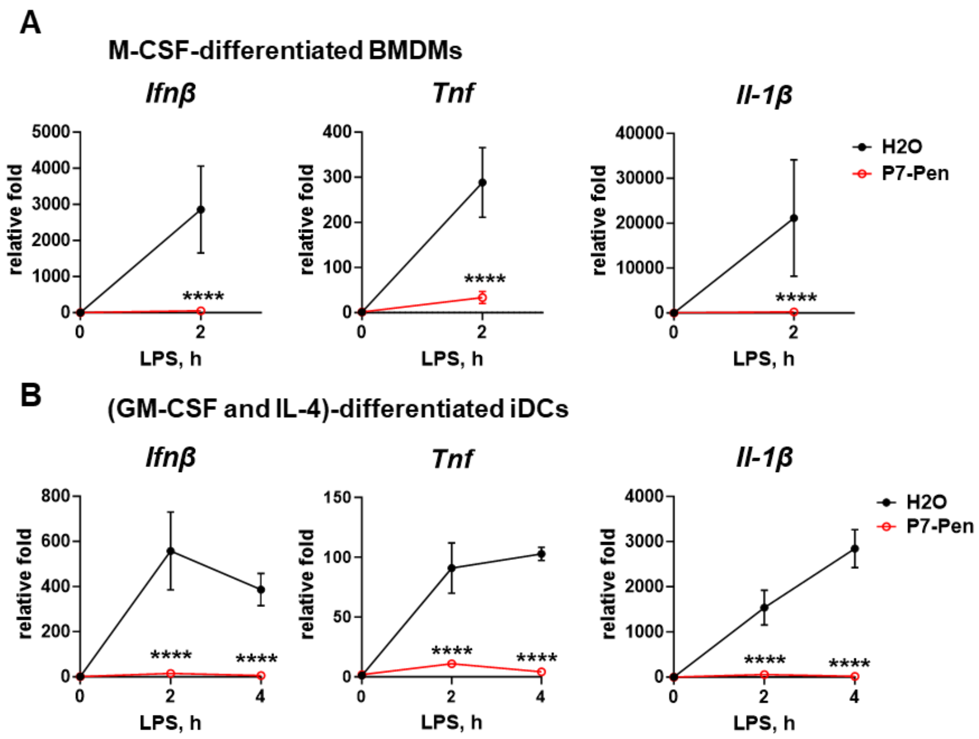


**Supplementary Figure 4. P7 inhibits LPS-mediated cytokine expression in primary murine BMDMs and BM-derived iDCs.** BMDMs (**A**) and iDCs (**B**) were generated from BM of C57BL/6 mice. (**A**) M-CSF-differentiated BMDMs and (**B**) BM-derived iDCs (GM-CSF and IL-4-differentiated) were pretreated with water solvent ($H_2O$) or 10 µM P7-Pen for 30 min and stimulated with LPS (100 ng/ml) for indicated time, along with R848 stimulations shown in the main Figure 5. Cells were analyzed by RT-qPCR for *Ifnβ, Tnf* and *Il-1β* mRNA expression. Data presented as relative fold change when compared to unstimulated sample pretreated by water ($H_2O$), mean ± SEM (n = 3). Statistical testing was done by two-way ANOVA on log-transformed data, significance levels: ****$P < 0.0001$.
